# Sub-3 Å resolution structure of 110 kDa nitrite reductase determined by 200 kV cryogenic electron microscopy

**DOI:** 10.1101/2020.07.12.199695

**Authors:** Naruhiko Adachi, Takahide Yamaguchi, Toshio Moriya, Masato Kawasaki, Kotaro Koiwai, Akira Shinoda, Yusuke Yamada, Fumiaki Yumoto, Takamitsu Kohzuma, Toshiya Senda

## Abstract

Cu-containing nitrite reductases (NiRs), a well-studied family of 110 kDa enzymes, play central roles in denitrification and have over 100 Protein Data Bank entries. However, such issues as crystal packing, photoreduction, and lack of high pH cases have impeded structural analysis of the catalytic mechanism. Here we show the cryogenic electron microscopy (cryo-EM) structures of *Achromobacter cycloclastes* NiR (*Ac*NiR) at 2.99 and 2.85 Å resolution with pH 6.2 and 8.1, respectively. Comprehensive comparisons with cryo-EM and 56 *Ac*NiR crystal structures suggested crystallographic artifacts in residues 185–215 and His255 due to packing and photoreduction, respectively. With electron paramagnetic resonance spectroscopy, a newly developed map comparison method supported local structural changes at pH 8.1 around the type-2 Cu site, including His255 deprotonation. While the theoretical coordination error estimation of cryo-EM structures remains difficult, combined analysis using X-ray and cryo-EM structures will allow deeper insight into the local structural changes of proteins.

## Introduction

Cu-containing nitrite reductases (hereafter, NiRs) (EC number 1.7.2.1), which catalyze the reduction of nitrite (NO_2_^−^) to nitric oxide (NO), play a key role in denitrification in bacteria^1^. The first NiR was discovered in *Alcaligenes xylosoxidans*^2^, and its orthologues were found in several bacteria such as *Achromobacter cycloclastes*^3^, *Alcaligenes faecalis*^4^, and *Rhodobacter sphaeroide*^5^. The NiRs are trimeric enzymes with a molecular mass of approximately 110 kDa. Each subunit contains two Cu ions, type-1 Cu (T1Cu) and type-2 Cu (T2Cu), and both are necessary for the enzymatic reaction. The T1Cu accepts an electron from pseudoazurin^6^ and donates the electron to the T2Cu^7^. Interestingly, the enzyme activity of NiRs shows high pH dependence with a bell-shaped activity curve. Crystallographic studies of several NiRs have been performed to understand the molecular mechanism of the catalytic reaction and its high pH dependence^8–13^. These crystal structures have revealed details of the T1Cu and T2Cu sites (**Supplementary Fig. 1**). The T1Cu is coordinated by His95, Cys136, His145, and Met150 (hereinafter, all residue numbers in this manuscript will refer to *Achromobacter cycloclastes* NiR (*Ac*NiR) unless otherwise specified). T2Cu, which is located at the interface of two subunits, is coordinated by His100, His135, His306’, and one water molecule (the prime mark at the end of the residue number indicates that the residue belongs to an adjacent subunit in the NiR trimer). The water (or hydroxide) coordinated to T2Cu is known to be replaced with nitrite.

The first intensive analysis for the pH dependence of NiRs was performed using *Ac*NiR, which has an optimal pH of 6.2. Although the crystal structures of *Ac*NiR under the pH range from 5.0 to 6.8 were determined, no significant structural changes were observed in this pH range^9^. Considering the optimal pH of *Ac*NiR, the pH range investigated was rather small for investigation of the pH effect on the enzyme activity. However, this was probably a limitation of the crystallographic method, because crystals are stable only in a limited pH range. Other interesting studies of the pH effect were performed using the NiR from *Rhodobacter sphaeroides* (*Rs*NiR)^12,14^. In those studies, the authors determined the crystal structures of *Rs*NiR at pH 6.0 and 8.4 and analyzed the structure around the T2Cu site. They revealed that the pH change induced a shift of the ligand water molecule and a slight conformation change of His255’. These crystallographic studies suggested that Asp98 and His255’ are involved in the catalytic reaction of NiRs^9,12,15^. Indeed, amino acid replacement of Asp98 and His255’ of NiRs resulted in a large reduction of the enzyme activities^16,17^. In addition to these structural studies, many biochemical and physicochemical analyses have been performed. Kinetic studies have supported a proton coupled electron transfer mechanism of NiRs^18,19^. Recently, an X-ray free electron laser (XFEL) study proposed that His255’ changes its conformation during the catalytic reaction, which could explain a redox coupled proton switch for proton coupled electron transfer^20^.

Although the reaction mechanism of NiRs has been explained based on their crystal structures, it should be noted that crystal packing can restrict intrinsic conformational changes and dynamics of the protein in the crystal. It is frequently the case that the crystal packing hampers the catalytic reaction in the crystal, resulting in the trapping of a reaction intermediate within the crystal^21^. Therefore, structural information without the crystal-packing effects should be included when analyzing the enzymatic mechanism. Single particle analysis (SPA) using cryogenic electron microscopy (cryo-EM) is a critical method for determining near-atomic resolution structures without a packing effect. Moreover, cryo-EM has the advantages that the condition of the sample environment, such as the pH, and salt concentrations, can be changed more easily than the crystallographic method.

With these advantages of the cryo-EM SPA, we determined the cryo-EM structures of *Ac*NiR at pH 6.2 and 8.1 to analyze its pH dependence and catalytic mechanism. A comprehensive comparison between the cryo-EM structures and crystal structures of *Ac*NiR revealed the packing effects on the crystal structures. In addition, the cryo-EM structures made it possible to examine pH-associated changes around the T2Cu site of *Ac*NiR.

## Results

### Optimization of parameters for SPA

For the SPA, 794 and 694 micrographs were collected at pH 6.2 and 8.1, respectively. Typical micrographs after the motion collection showed that ice-embedded *Ac*NiR was monodisperse (**Fig. 1a, b**). Particles of *Ac*NiR were picked up and subjected to two-dimensional (2D) classification (**Fig. 1a, b**). The orientation distributions were uniform in both cases (**Supplementary Fig. 2a, c**). The statistics of data collection, map reconstruction, and model refinements of the cryo-EM SPA for *Ac*NiR are summarized in **Supplementary Table 1**.

**Fig. 1.**
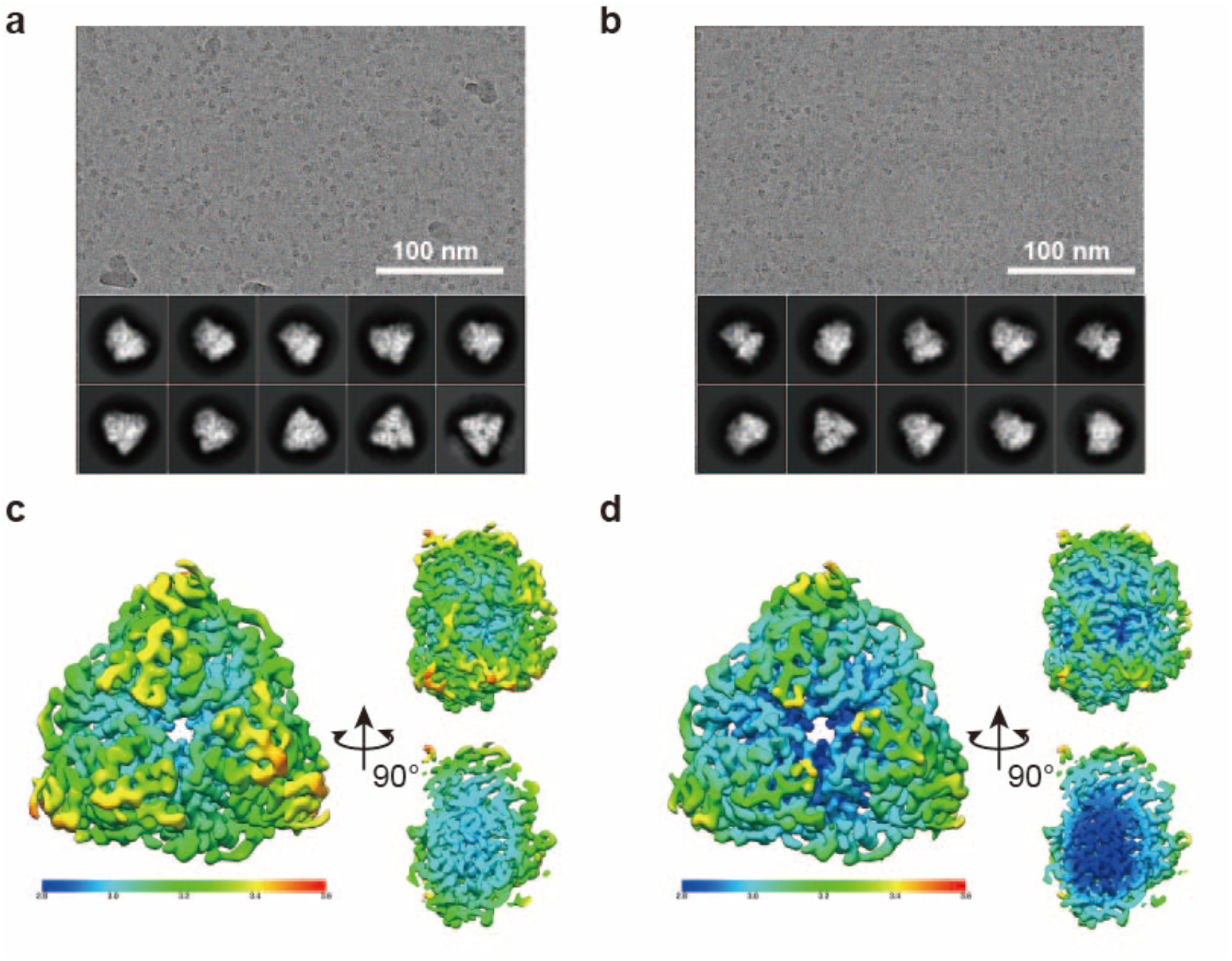
Cryo-EM SPA of *Ac* NiR at pH 6.2 and 8.1. **a, b** Typical micrographs and two-dimensional class averages of cryo-EM for *Ac*NiR at pH 6.2 (a) and 8.1 (b). **c, d** Local resolution of cryo-EM structures of *Ac*NiR at pH 6.2 (c) and 8.1 (d).

The box sizes and particle mask diameters were carefully optimized to achieve resolution better than 3.0 Å for *Ac*NiR, the molecular mass of which was approximately 110 kDa. Theoretically, the optimal box size for a given defocus range should eliminate the CTF aliasing artifacts. Unlike that for large proteins, the choice of box size for small proteins is not dominated by the true size of the particles but rather by the CTF aliasing frequency limit^22^. This is because the limit is unrelated to the particle size but mainly depends on the defocus steps used in the cryo-EM session. Furthermore, the optimal particle mask diameter should enhance the restoration of information by CTF correction in the 3D refinement step. The particle mask diameter depends on not only the true size of the particle, but also the width of the point spread function (PSF), which, again, mainly depends on the defocus value. In real space, the information of the projected particle view is spread outside of the true particle edge by PSF. Ideally, the particle mask diameter should enclose the information that is spread outside of the particle edge by the PSF^23^.

For the *Ac*NiR datasets under both the pH 6.2 and pH 8.1 conditions, the optimal box size was 486 pixels at the original pixel size of 0.88 Å/pixel. This ensured that there was no CTF aliasing up to 2.62 Å for a 2.5 μm defocus, which was the maximum defocus value of both datasets used in the final stage of the processing. To check whether this box size was truly optimal, the three cycles of CTF refinement and Bayesian polishing steps were repeated with two additional box sizes using the pH 8.1 dataset (see also Methods). The box sizes of 486, 384, and 256 pixels resulted in final resolutions of 2.87, 2.96, and 3.08 Å, respectively, supporting the choice of the 486-pixel box size as optimal. To our knowledge, the conventional criterion states that the box size should be 1.5–2.0 times the longest particle view. Since the measured diameter of the *Ac*NiR particle was ~100 Å (~114 pixels with 0.88 Å/pixel), the convention suggests that the box size should be 256 pixels or even smaller. That is, the box size optimization alone yielded a resolution improvement of 0.21 Å (from 3.08 to 2.87 Å) with this dataset.

Next, the optimal particle mask diameter was selected by executing multiple runs of initial 3D refinement right after scaling up the pixel size back to the original 0.88 Å/pixel (see also Methods) and choosing the diameter that achieved the highest resolution. With the pH 8.1 *Ac*NiR dataset, the particle mask diameters of 100, 132, and 164 Å were used and resolutions of 3.62, 3.39, and 3.34 Å were obtained, respectively. 164 Å was selected as the optimal particle mask diameter. Again, the measured *Ac*NiR particle diameter was ~100 Å. Therefore, the resolution improvement due to the optimization of particle mask diameter was about 0.28 Å (from 3.62 to 3.34 Å). In total, the optimization of the box size and particle mask diameter improved to approximately 0.49 Å for the pH 8.1 *Ac*NiR reconstruction compared with the widely used criteria. Using the above optimal parameter values, the cryo-EM maps at the highest resolution were obtained. Their resolutions were estimated as 2.99 Å resolution at pH 6.2 and 2.85 Å resolution at pH 8.1 based on the criterion of 0.143 Fourier Shell Correlation (FSC) (**Supplementary Fig. 2b, d**)^24^. Local resolution was also calculated (**Fig. 1c, d**).

Before model building, the map quality was examined by the d_99_ value using *phenix.mtriage*. The highest resolution Fourier map coefficients may or may not contribute significantly to the map. The d_99_ value represents how many of the highest resolution coefficients can be omitted before the corresponding real-space map changes significantly^25^. The d_99_ values of the cryo-EM maps at pH 6.2 and 8.1 were 3.02 and 2.89 Å, respectively. These were nearly the same as the resolutions from FSC plots, suggesting that the obtained maps were of reasonable quality. The scattering potential maps showed the triangle prism shape of the *Ac*NiR molecule, which is consistent with the small angle X-ray/neutron scattering data from the previous reports^26,27^. The correlation coefficients of the final model and the masked map for the macromolecule only (CC_mask_) at pH 6.2 and 8.1 were 0.88 and 0.87, respectively.

### Overall structure comparison between the cryo-EM and crystal structures

The cryo-EM structures have the same fold as obtained from X-ray crystallography. The least-squares (LSQ) fitting of the two cryo-EM structures using 321 CA atoms (residues 10–330) gave a root-mean-square XYZ displacement (RMSD) of 0.281 Å, suggesting that no large conformational changes were associated with a pH change from 6.2 to 8.1. On the basis of this fact, the cryo-EM structure determined at pH 6.2 was used for further analysis unless stated otherwise.

To analyze the packing effect on the crystal structures of *Ac*NiR, we compared the crystal structures of *Ac*NiR in the Protein Data Bank (PDB) with the cryo-EM structure (pH 6.2). Initially, a comparison was made using the cryo-EM structure and a 0.9 Å-resolution crystal structure (PDB ID: 2bw4) that was used as an initial model for the model building. An LSQ fitting of the two structures gave an RMSD value of 0.697 Å. This RMSD value was significantly larger than the averaged RMSD (0.36 ± 0.17 Å) obtained from the pairwise LSQ fittings of all *Ac*NiR crystal structures (56 structures) in PDB (**Supplementary Tables 2 and 3**), suggesting that the cryo-EM structure has significant deviations from the crystal structures. We considered that the difference of the two RMSD values may have arisen from the packing effect of the crystal structures, and thus further comparisons between the cryo-EM and crystal structures of *Ac*NiR were performed. Interestingly, comprehensive comparison of the cryo-EM structure with all *Ac*NiR structures in PDB revealed that the *Ac*NiR crystal structures in PDB can be classified into two distinct groups, A and B (**Fig. 2a, Supplementary Table 2**). 1nia^9^ and 6gsq^28^ appear to represent characteristics of groups A and B, respectively (**Supplementary Fig. 3**). Although the crystal structures in groups A and B are very similar to each other, significant structural differences were found in residues 185–215. The conformations of residues 185–215 of the cryo-EM structure are similar to those in group A (**Fig. 2b**). The conformational difference of this region between the cryo-EM and X-ray structures of group B was confirmed by a 3D density existence probability map that was devised in this study to calculate quantitative significance of this local structural difference (**Fig. 2c, d**).

**Fig. 2.**
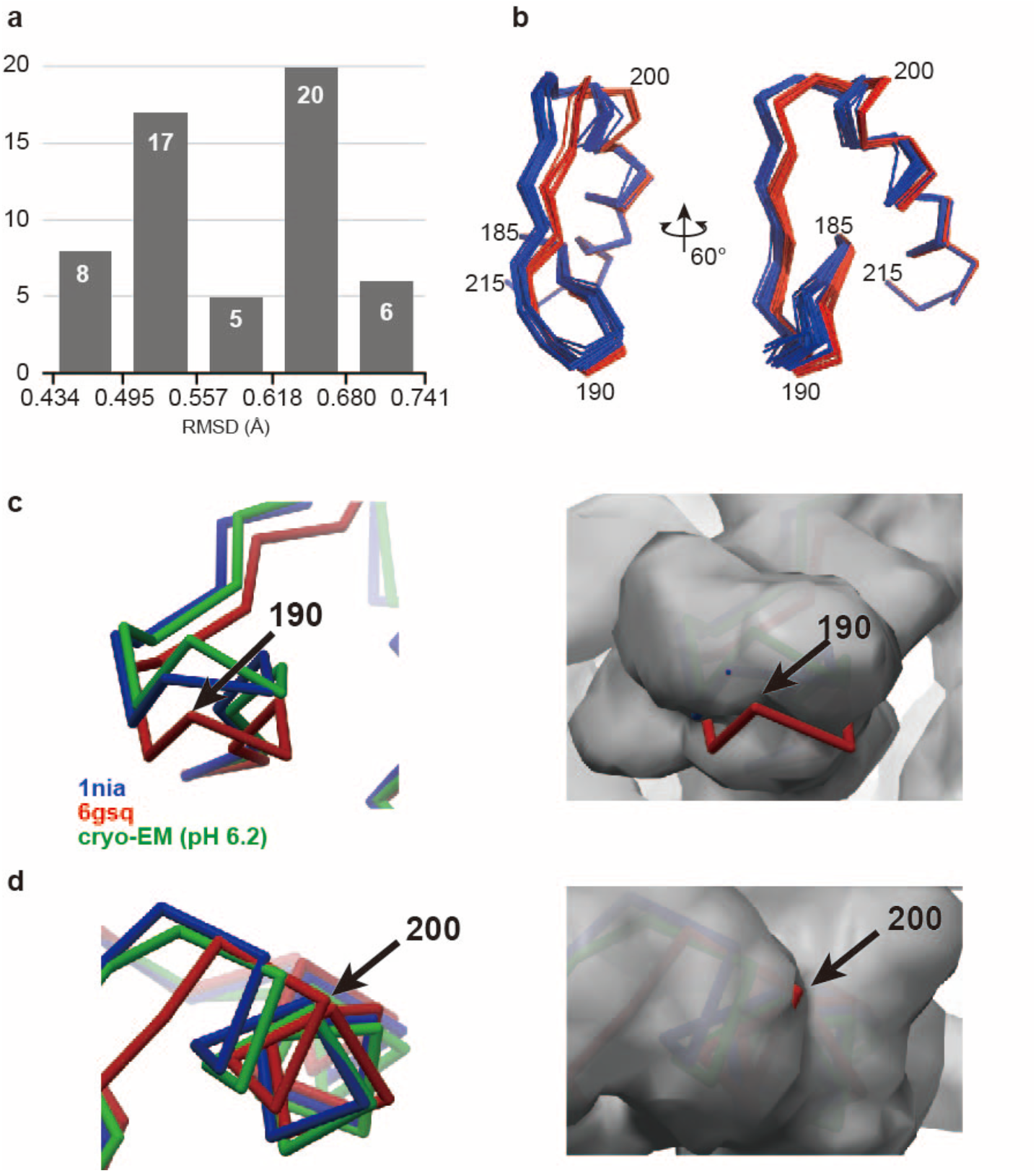
Two conformations in residues 185-215. **a** Histogram showing the distribution of RMSD values of 56 *Ac* NiR crystal structures in PDB with respect to the cryo-EM structure (pH 6.2) of *Ac* NiR. **b** Superposition of 56 PDB structures of residues 185–215. The structures of group A and B (Supplementary Table 2) are shown in blue and red, respectively. **c, d** Superposition of 1nia (blue), 6gsq (red), cryo-EM structures (left) and the 3D probability map of the corresponding region (right). The contour level is 0.125, meaning that a voxel value lower than 1/8 indicates no probability of any density existence at this 3D position with a 7/8 confidence level. Panels **c** and **d** show the conformational differences around residues 190 and 200, respectively.

While residues 185–215 showed the largest difference between the cryo-EM and X-ray structures, structural superposition revealed small but significant structural differences in other regions. To analyze these differences in a quantitative manner, the averaged displacement for each CA atom was calculated using the LSQ-fittings of crystal structures onto the cryo-EM structure (pH 6.2). Although the averaged displacements of CA atoms were large near the surface of the trimer, the averaged displacements were relatively small around the T2Cu site (**Fig. 3a, b**). Another interesting observation was the relationship between the unit-cell parameters and the RMSD values of the crystal and cryo-EM structures. Most of the crystal structures belonged to the cubic space group *P*2_1_3. The RMSD value showed a strong correlation with the length of the *a*-axis of the cubic crystal lattice (**Fig. 3c**).

**Fig. 3.**
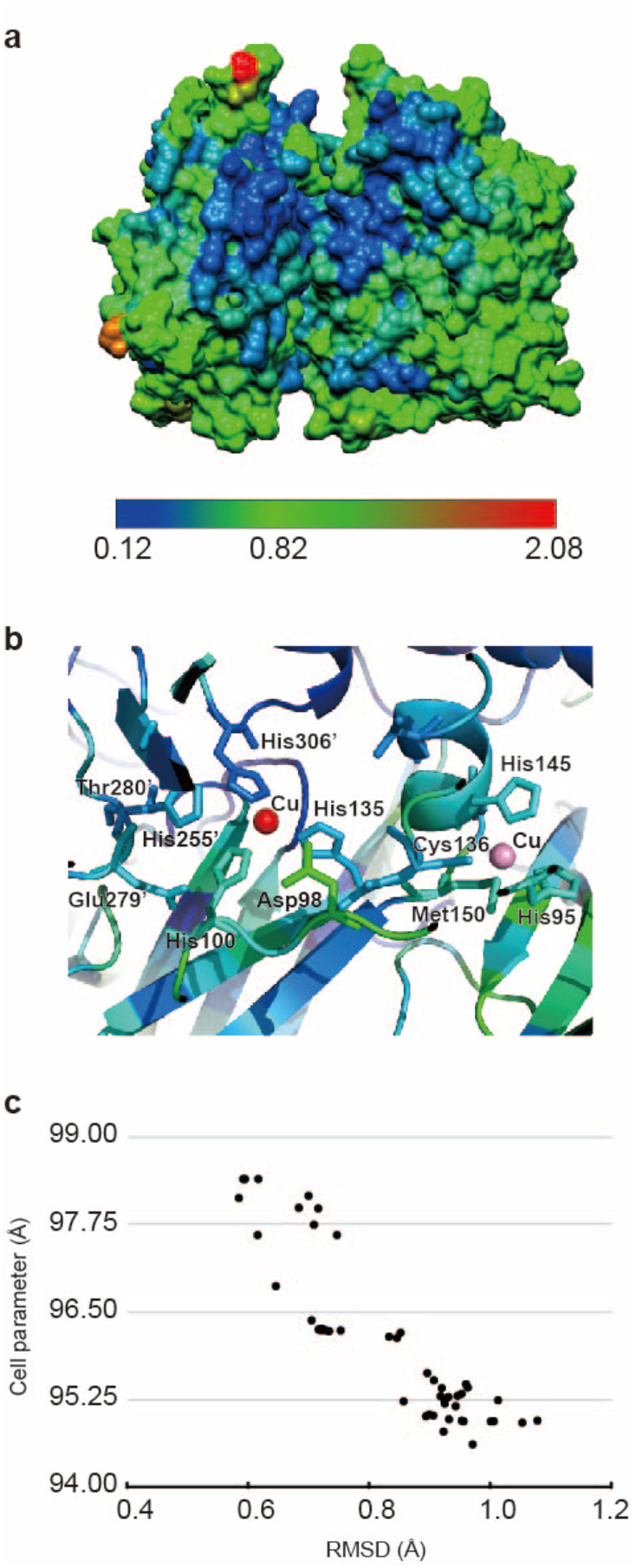
Packing effects of the crystal structures of *Ac* NiR. **a** Distribution of the averaged displacements of CA atoms shown on a half-cut structure of *Ac* NiR. The magnitudes of the displacements are shown with rainbow colors, from blue to red, as shown in the color bar below. **b** Distribution of the averaged displacements around the T1Cu and T2Cu sites. Molecules are colored in the same manner as in panel **a**. **c** Correlations between the RMSD values calculated from the crystal structures and corresponding unit cell parameters of the crystals. The RMSD values were obtained from the LSQ fitting of the cryo-EM structure (pH 6.2) and *Ac*NiR crystal structures of the cubic space group *P*2_1_3.

### Comparison of the coordination sphere between the cryo-EM and crystal structures

Next, we examined local differences between the cryo-EM (both pH 6.2 and 8.1) and X-ray structures, particularly around the T2Cu coordination sphere, where the effects of the crystal packing were relatively small. The coordination distances of the cryo-EM structures showed only small differences, approximately 0.1 Å, from those of the crystal structures (**Table 1**). LSQ-fittings of the crystal structures onto the cryo-EM structure (pH 6.2) showed a positional shift by approximately 0.6 Å of His100, which may have been a packing effect. However, no significant difference of the coordination distances was found between His100 and the Cu ion at both the pH 6.2 and pH 8.1 cryo-EM structures (**Table 1**). It is of note that the T2Cu has one water (or hydroxide) ligand in addition to the His ligands in the crystal structures of the substrate-free form. While no clear cryo-EM density for the ligand water molecule was observed at the T2Cu site, we found a protrusion of the cryo-EM density in the direction of the water ligand in the crystal structures in both the pH 6.2 and 8.1 cryo-EM structures (**Fig. 4a**). Considering the resolutions of the cryo-EM structures, this protruded cryo-EM density could be interpreted as a ligand water molecule; it seems possible that a water molecule could be placed at the vacant position of the tetrahedral coordination sphere. The position of the possible water ligand was nearly the same between the pH 6.2 and 8.1 structures.

**Table 1.**
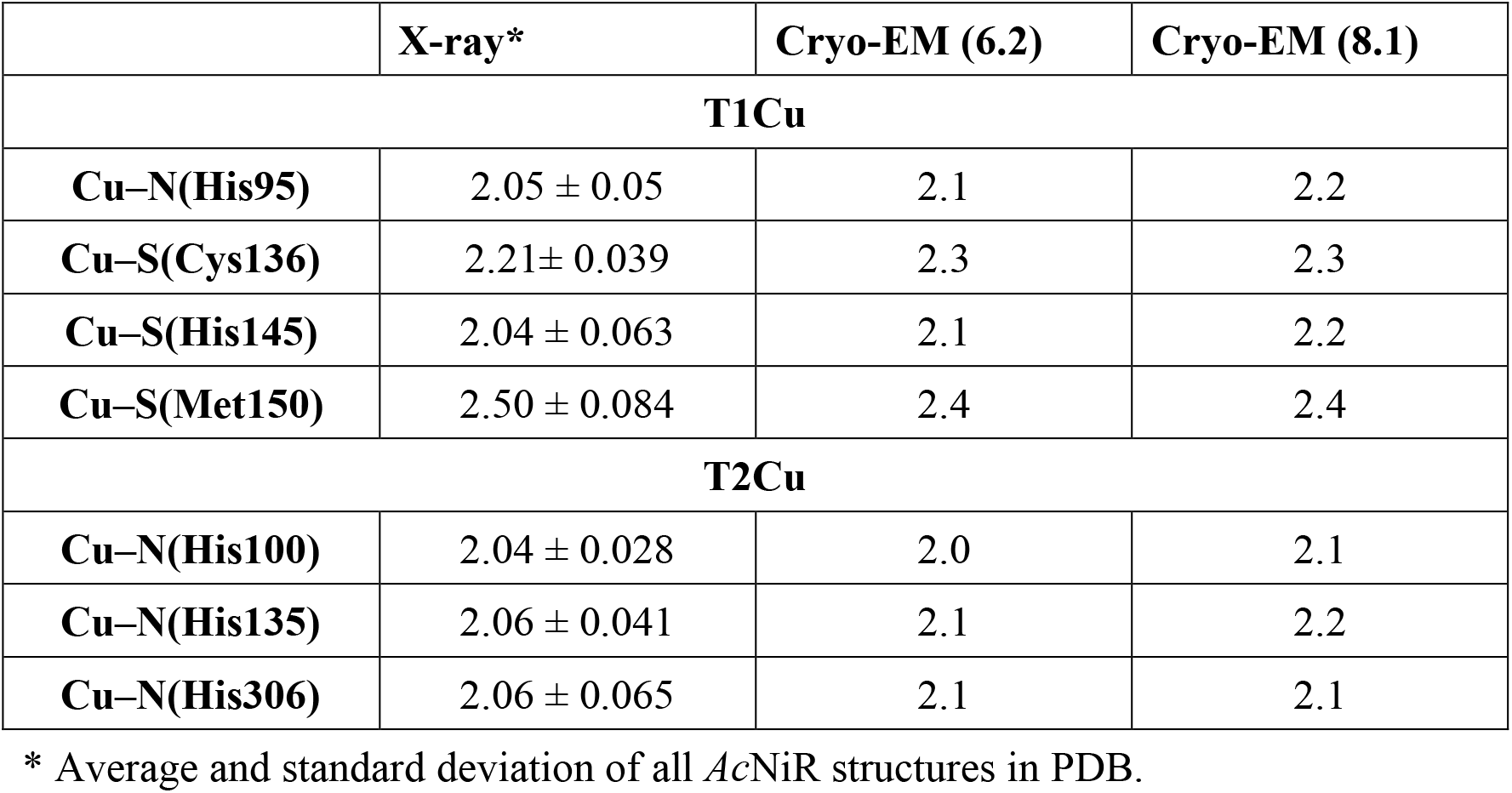
Coordination sphere geometries of the T1Cu and T2Cu sites of *Ac* NiR.

|  | **X-ray\*** | **Cryo-EM (6.2)** | **Cryo-EM (8.1)** |
| --- | --- | --- | --- |
| **T1Cu** |  |  |  |
| **Cu–N(His95)** | 2.05 ± 0.05 | 2.1 | 2.2 |
| **Cu–S(Cys136)** | 2.21± 0.039 | 2.3 | 2.3 |
| **Cu–S(His145)** | 2.04 ± 0.063 | 2.1 | 2.2 |
| **Cu–S(Met150)** | 2.50 ± 0.084 | 2.4 | 2.4 |
| **T2Cu** |  |  |  |
| **Cu–N(His100)** | 2.04 ± 0.028 | 2.0 | 2.1 |
| **Cu–N(His135)** | 2.06 ± 0.041 | 2.1 | 2.2 |
| **Cu–N(His306)** | 2.06 ± 0.065 | 2.1 | 2.1 |

\* Average and standard deviation of all *AcNiR* structures in PDB.

**Fig. 4.**
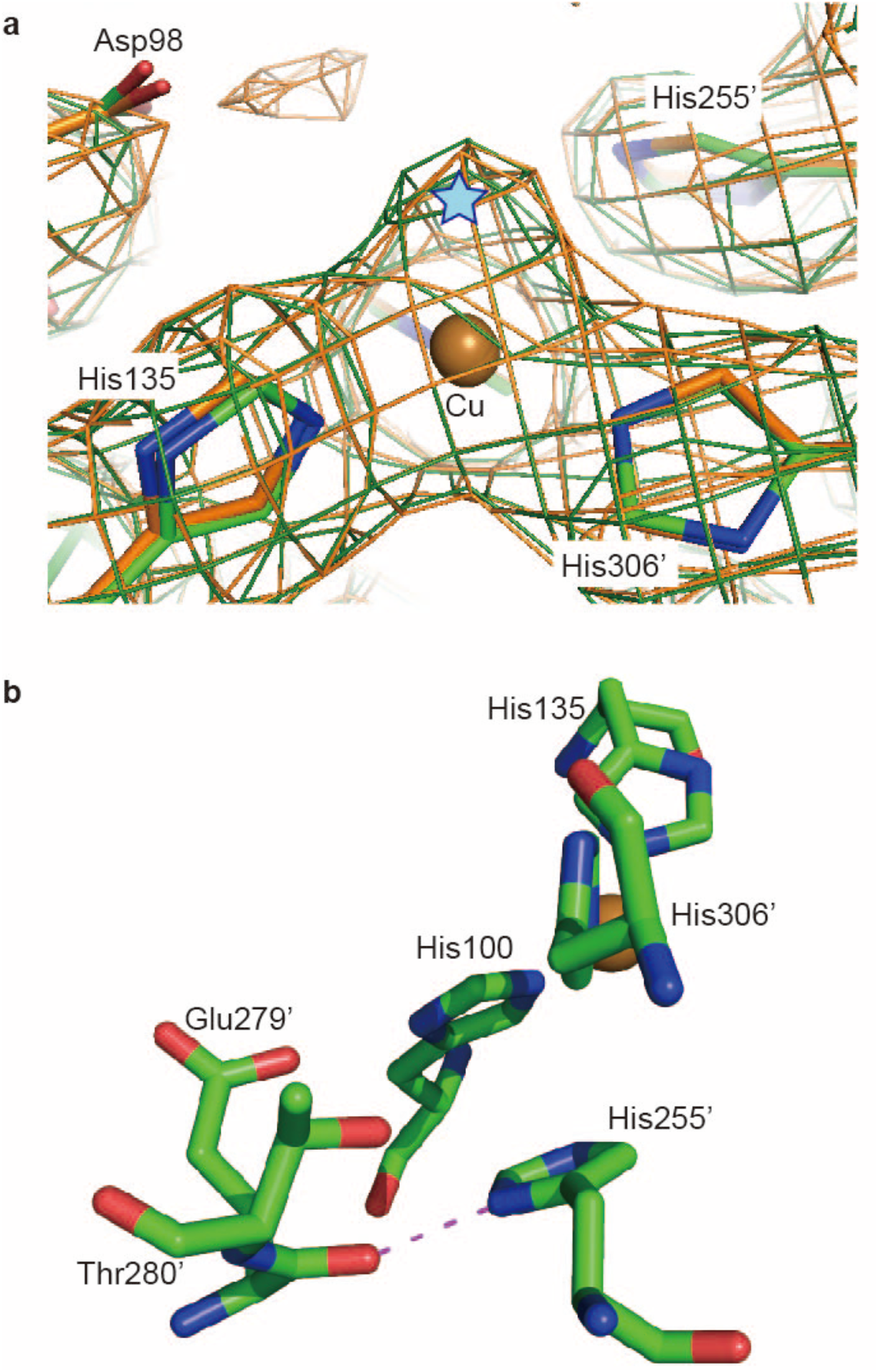
Structure around the T2Cu site. **a** Cryo-EM densities around the T2Cu site at pH 6.2 (green mesh) and 8.1 (orange mesh). CA atoms of the cryo-EM structures at pH 6.2 and 8.1 are shown in green and orange sticks, respectively. The possible position of the ligand water molecule is indicated by a cyan star. **b** A hydrogen bond between His255’ and Glu279’ is indicated by a purple dotted line.

In the outer coordination sphere of *Ac*NiR, the conformation of His255’ has been intensively studied due to its involvement in the catalytic reaction^11,13,16,20,29^. Interestingly, this residue showed conformational variety among the crystal structures due to photoreduction by X-ray^12,14,20^. The crystal structures in PDB showed three types of interactions of the ND atom of His255’: the ND atom forms a hydrogen bond with (1) the OG atom of Thr280’, (2) the main-chain carbonyl oxygen of Glu279’, and (3) both of the OG atom of Thr280’ and the carbonyl oxygen of Glu279’ weakly (**Supplementary Fig. 4**). In both cryo-EM structures (pH 6.2 and 8.1), the ND atom of His255’ forms a hydrogen bond with the main chain carbonyl oxygen of Glu279’ with a distance of 2.46Å (**Fig 4b**). This conformation of His255’ suggests that the cryo-EM structures of *Ac*NiR represent the oxidized form^20,30^.

### Comparison of the T2Cu sites between the pH 6.2 and 8.1 conditions

Next, we examined whether or not pH-associated local conformation changes exist around the T2Cu site of the cryo-EM structures. Although the two cryo-EM structures are essentially the same, small differences were found in the outer coordination sphere.

A comparison of the two structures revealed that the direction of the C-O bond of the main chain carbonyl oxygen of Glu279’ at pH 8.1 is different from that at pH 6.2, resulting in an approximately 0.7 Å shift of the carbonyl oxygen. This difference, in turn, seemed to affect the χ2 angle of His255’, which was altered by approximately 5 degrees (**Fig. 5**). However, since these changes were relatively small, we developed a new method, the equal-volume 3D difference map method, to confirm these differences. This method visualizes differences between two maps. However, since typical cryo-EM maps have arbitrary scales, two maps should be normalized before subtraction. In this study, two maps were normalized by generating the corresponding binarized maps. The binarized maps have only two values for each grid, 1 and 0, which correspond to the regions inside and outside the *Ac*NiR structure in the cryo-EM density map, respectively. To prepare the binarized map, we first estimated a contour level for each cryo-EM map in order to make the (protein) volumes of the two maps nearly identical. Importantly, the binary map should express the shapes of the side chains well. Then, the contour level of each binary map was optimized to make the volume of the two maps exactly the same. After obtaining the normalized binary maps, one of the binarized maps was subtracted from the other to make an equal-volume 3D difference map. As shown in **Fig. 5**, there is a pair of difference densities around the carbonyl oxygen of Glu279’ in the equal-volume 3D difference map (blue and red blobs labeled “A”). In addition, the small change of the side chain position of His255’ seems to be supported by the map (blue and red blobs labeled “B”). However, the method seems to be sensitive to the threshold value used in the binarization step (**Supplementary Fig. 5**) and so it should be considered as a guide or support rather than a stand-alone proof. The interpretation of the difference blobs was done carefully with other types of measurements.

**Fig. 5.**
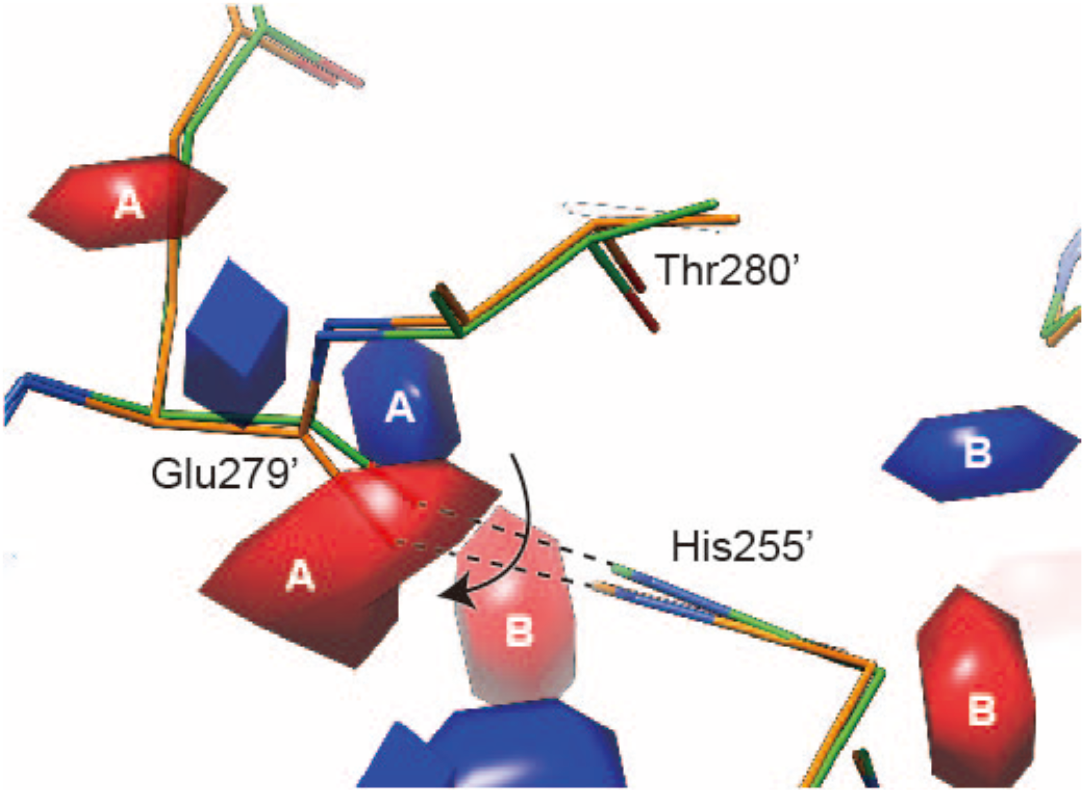
Conformational changes of His255’ and Glu279’ associated with increase of pH from 6.2 to 8.1. Conformational changes of His255’ and Glu279’. CA atoms of cryo-EM structures at pH 6.2 and 8.1 are shown in green and orange, respectively. Interactions between His255’ and mainchain carboxyl oxygen of Glu279’ are indicated by the dotted lines and the direction of change from pH 6.2 to 8.1 by the allow. The equal-volume 3D difference blobs between the pH 6.2 and pH 8.1 cryo-EM density maps are also shown. The blue and red blobs are densities existing only in the pH 6.2 and pH 8.1 maps, respectively. The blobs which assumed to be associated to Glu279’ and His255’ are labeled as “A” and “B”, respectively.

### Electron paramagnetic resonance (EPR) spectroscopy

The cryo-EM structures of *Ac*NiR at pH 6.2 and 8.1 demonstrated no significant structural differences around the T2Cu site other than small changes around His255’. To obtain detailed information around the T2Cu site, including information on the deprotonation, the X-band EPR spectra of *Ac*NiR at pH 6.0 and 8.0 were measured (**Fig. 6**). The T1Cu and T2Cu sites in the oxidized form are paramagnetic and detectable by EPR^10,12,31,32^. The EPR spectra of *Ac*NiR at pH 6.0 and 8.0 showed significant differences (**Supplementary Table 5**). The g_‖_ and A_‖_ values were obtained as 2.18 and 7.2 mT for the T1Cu site at pH 6.0 by simulation, and these values are consistent with the parameters in the earlier report (g_‖_ = 2.18 and A_‖_ = 7.2 mT)^7^. The simulated parameters of the T1Cu site at pH 8.0 were identical to the values at pH 6.0, indicating that the Cu environment at the T1Cu site was not affected by alkaline pH change. In contrast, the EPR spectra of the T2Cu site showed differences between pH 6.0 and 8.0. The EPR spectrum of the T2Cu site at pH 6.0 was a single component, and the g_‖_ and A_‖_ values were 2.34 and 13.1 mT, respectively. At pH 8.0, the T2Cu EPR signal in the g_‖_ region was changed by the appearance of an additional signal (g_‖_ = 2.27 and A_‖_ = 16.2 mT). This suggests that the T2Cu site structure was affected by the protonation/deprotonation of the adjacent amino acid(s). The different EPR spectroscopic features at the T2Cu site reflect the structural change of His255 observed on the cryo-EM map of *Ac*NiR. The EPR spectra of NiR from *Alcaligenes faecalis* (*Af*NiR) studied by Boulanger et al. showed that the g_‖_ value of the T2Cu site increased by 0.20 due to the His255Asn mutation^16^. This fact supports that the changes observed around the T2Cu site were due to the protonation/deprotonation of His255’.

**Fig. 6.**
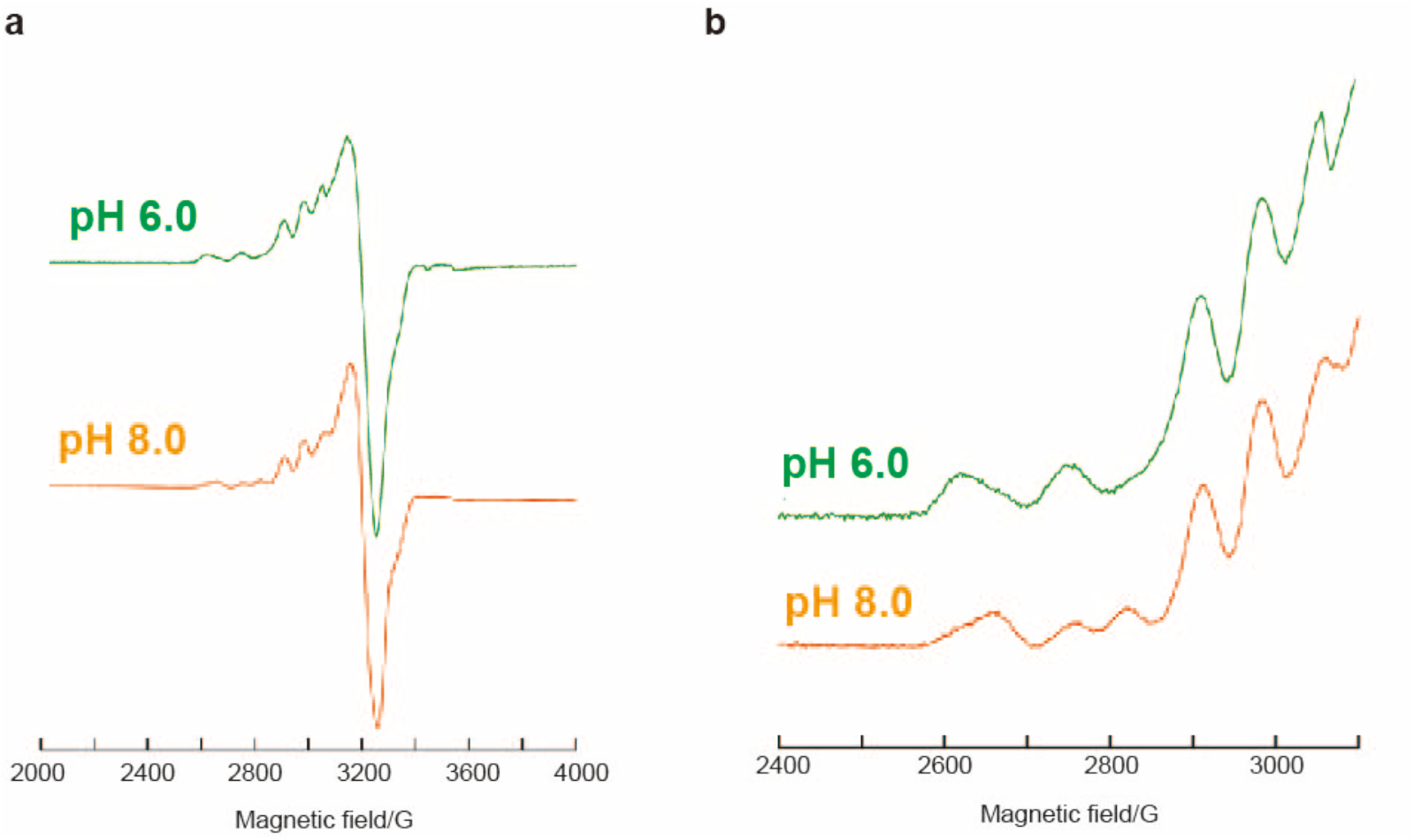
EPR spectra of AcNiR. **a** X-band EPR spectra of AcNiR at pH 6.0 and 8.0, and **b** the A_‖_ region of these spectra.

## Discussion

NiRs are among the most well-studied enzymes and have been analyzed by biochemical, crystallographic, spectroscopic, and computational methods. Most of the 3D structural information of NiRs has been obtained by X-ray crystallographic analyses, which have revealed the reaction intermediates of NiRs, the binding geometry of nitrite, and other critical structural information^8–13,15^. However, the crystal structure of NiRs is known to be affected by crystal packing, and sometimes packing effects modulate the enzyme properties in crystals^21,33^. Therefore, 3D structures without packing effects should be considered when analyzing the catalytic reaction of enzymes with crystal structures. Furthermore, other artifacts in the crystal structure, such as the pH environment and photoreduction, should be corrected by other methods. While several methods have been used to compensate for these artifacts, cryo-EM SPA is becoming a powerful candidate that can complement the disadvantages of the crystallographic method. For this purpose, however, it is necessary to (1) obtain sufficiently high resolution cryo-EM structures, (2) measure the structure reliabilities, and (3) analyze their structure differences. In this study, we investigated pH-associated structural differences of NiRs in an effort to satisfy all three requirements.

We succeeded in determining the cryo-EM structures (pH 6.2 and pH 8.1) of *Ac*NiR at better than 3 Å resolution; among the cryo-EM structures at better than 3 Å resolutions using conventional 200 kV instrument, *Ac*NiR has one of the smallest molecular sizes. Our analysis demonstrated that optimization of the SPA parameters, specifically the box size and particle mask diameter, was critically important to obtain the high-resolution structures; for a given defocus distribution of a dataset, the optimal values for the box size and particle mask diameter are different between 200 kV and 300 kV electron microscopes because the different electron wavelengths yield different contrast transfer functions (CTFs). It should be noted that the cryo-EM SPA method allowed us to determine the structure of *Ac*NiR at pH 8.1 for the first time, because it was much easier to change the pH value of the sample by the cryo-EM SPA method than by the crystallographic method.

The RMSD value for the fitting of the two cryo-EM structures under pH 6.2 and 8.1 was 0.28 Å, which is marginally smaller than the averaged RMSD value derived from comprehensive pairwise LSQ fittings of the 56 *Ac*NiR crystal structures in PDB, but not significantly different (0.36 ± 0.17 Å; **Supplementary Tables 3**). The small RMSD value suggests the good reproducibility and accuracy of the two cryo-EM structures obtained in this study. Since there are no conventional measures to estimate coordinate errors of the cryo-EM structures, it is difficult to assess the significance of the difference between the two cryo-EM structures. At the same time, however, comprehensive comparison of the cryo-EM structures with the crystal structures yielded no evidence that these cryo-EM structures contained significantly larger coordinate errors than those of the crystal structures. Analysis of the packing effect through a comparison of the cryo-EM and crystal structures revealed that the conformations of the residues near and on the surface of the molecule were more affected by the crystal packing than those located inside the molecule. This result is physically reasonable and, in turn, seems to support the accuracy of the cryo-EM structures obtained in this study.

The largest packing effect of the crystal structure was found in the residue range from 185 to 215. The structural difference of this region is obvious, and this difference was quantitively confirmed by the 3D density existence probability map that we devised in this study (**Fig. 2c, d**). To our knowledge, this is one of the first attempts to establish a quantitative measure to evaluate the confidence level of density existence at each 3D position in a cryo-EM structure without the necessity of modeling. Comparative analysis of the *Ac*NiR structures suggested that the conformations of this region in the crystal structures were affected by physical contacts such as crystal packing (**Fig. 3c**). When the packing effect is large—i.e., when the cell parameter (length of the *a* axis) is small—the conformation of this region is more deviated from that of the cryo-EM structures. However, when the packing effect becomes relatively small by increasing the cell parameter, the conformation of this region changes in the crystal, more closely resembling that of the cryo-EM structure. This fact suggests that residues 185–215 of *Ac*NiR in solution have conformations similar to those of the cryo-EM. Since this region is expected to be a part of the pseudoazurin-binding site^34^, the two conformers could play a regulatory role to form a binding site of pseudoazurin. There are some examples showing that the conformation change is related to the redox-dependent affinity control between electron transfer proteins^33,35–38^.

The oxidized-form conformation of His255 observed in the cryo-EM structures shows another advantage of the cryo-EM SPA. As reported earlier, many crystal structures of NiRs have suffered from photoreduction, resulting in the reduced form structure of His255 (**Supplementary Fig. 4**). Since His255 is a critical residue in the catalytic reaction, its conformation should be determined with great care. Serial femtosecond crystallographic analyses using XFEL and neutron diffractions have been utilized to avoid radiation damage on the crystal, and these approaches have given the oxidized-form structures of His255 in NiRs. However, neither method can avoid the packing effects. Therefore, high resolution cryo-EM structures could play a critical role in the study of redox enzymes, by freeing researchers from the longstanding struggles against packing and radiation damage to redox enzymes in the crystallographic method.

On the basis of the cryo-EM structures, we analyzed the characteristics of the T2Cu site of *Ac*NiR. The tetrahedral coordination sphere with three His residues and one water molecule, which has been demonstrated by the crystal structures, seemed to be conserved in the cryo-EM structures (**Fig. 4a**). *Rs*NiR showed a pH-dependent shift of the ligand water molecule in the T2Cu coordination sphere^12^. A similar water shift associated with pH change was also observed in the NiR derived from *Alcaligenes xylosoxidans*^39,40^. However, the cryo-EM structures in our study gave no evidence supporting a pH-associated shift of the ligand water molecule at the T2Cu site. Since many crystal structures of NiRs seem to be photo-reduced by synchrotron X-ray radiation^20,30^, it may be that the observed shift in the ligand water molecule was caused by photoreduction and/or packing effects.

In this study, we determined the near-atomic resolution structures of *Ac*NiR under two different conditions using a newly developed method for SPA parameter optimization. The cryo-EM SPA allowed us to compare the structures at pH 6.2 and 8.1. Moreover, a comprehensive comparative study with earlier determined crystal structures of *Ac*NiR revealed some changes associated with pH and physical interactions. While theoretical estimation of the coordination error of the cryo-EM structures remains difficult, by combining X-ray and cryo-EM structure analysis it becomes possible to study the local structure changes of proteins.

## Methods

### Preparation of nitrite reductase

Nitrite reductase was purified from the nitrate cultivation of *Achromobacter cycloclastes*. The cultivated bacteria in aqueous suspension were homogenized by ultrasonication, and the crude proteins in the soluble fraction were applied in cation-exchange and size-exclusion chromatography as described in a previous report^3^. The purity of *Ac*NiR was confirmed by sodium dodecyl sulfate poly acrylamide gel electrophoresis (SDS-PAGE). The samples for the cryo-EM experiments were prepared in the 100 mM potassium phosphate buffer at pH 6.2 and 8.1. The concentrations of *Ac*NiR were ca. 10-15 μM.

### Cryo-EM sample preparation and data acquisition

3 μL of sample was applied onto a holey carbon grid (Quantifoil, Cu, R1.2/1.3, 300 mesh) rendered hydrophilic by a 30 s glow-discharge in air (11 mA current with PIB-10). The grid was blotted for 15 s (blot force 0) and flash-frozen in liquid ethane using Vitrobot Mark IV (FEI) at 18 °C and 100% humidity.

Images were acquired with a Talos Arctica (FEI) microscope operating at 200 kV in the nanoprobe mode using the EPU software for automated data collection. For both the pH 6.2 dataset and pH 8.1 dataset, the movie frames were collected by a 4k × 4k Falcon 3 direct electron detector (DED) in electron counting mode at a nominal magnification of 120,000, which yielded a pixel size of 0.88 Å/pixel. Forty-nine movie frames were recorded at an exposure of 1.02 electrons per Å^2^ per frame, corresponding to a total exposure of 50 e^−^/Å^2^. The defocus steps used were 1.5, 2.0, 2.5, and 3.0 μm at pH 6.2, and 1.0, 1.5, 2.0, 2.5, and 3.0 μm at pH8.1.

### Cryo-EM data processing

Movie frames were aligned, dose-weighted, and averaged using MotionCor2^41^ on 5 × 5 tiled frames with a B-factor of 300 applied in order to correct for beam-induced specimen motion and to account for radiation damage by applying an exposure-dependent filter. The micrographs whose total accumulated motions were larger than 1500 Å were discarded. The non-weighted movie sums were used for CTF estimation with the program Gctf^42^ (512-pixel box size, 30 Å minimum resolution, 4 Å maximum resolution, 0.10 amplitude contrast), while the dose-weighted sums were used for all subsequent steps of image processing. The images whose CTF max resolutions were better than 6.0 Å were selected.

For *Ac*NiR at pH 6.2, the particles were picked up using SPHIRE crYOLO with a generalized model^43^ using a box size of 114 pixels and selection threshold of 0.3. The micrographs that contained less than 30 picks were excluded. A stack of 285,529 particle images was extracted from 713 dose-weighted sum micrographs while rescaling to 3.3 Å/pixel with 64-pixel box size, and subjected to reference-free 2D classification (100 expected classes, 120 Å mask diameter) in RELION3^44^. The 284,810 particles corresponding to the best 16 classes that displayed secondary-structural elements and multiple views of *Ac*NiR were selected for RELION3 *ab initio* reconstruction (asymmetry, single expected class, 100 Å mask diameter). C3 symmetry was imposed on the generated volume, and then the volume was low-pass filtered to 25 Å and used as an initial model for RELION3 3D classification (4 expected classes). One out of the four 3D classes clearly consisted of bad images containing non-targeted objects and was removed from the dataset. The volume of the best 3D class was rescaled to 0.88 Å/pixel with 486-pixel box size, low-pass filtered to 15 Å, and used as an initial model for the subsequent 3D refinement. Accordingly, 166,187 particles were also re-centered and re-extracted using the same rescale settings, and 3D auto-refined (C3 symmetry, 136 Å mask diameter, no padding) with a soft-edged 3D mask (no pixel extension, 6-pixel soft cosine edge) using RELION3. Prior to the particle extraction, the micrographs with defocus of >2.5 um were excluded to keep the box size at a practical level. In addition, micrographs showing obvious signs of ice crystallization (i.e., a strong ice ring in the Fourier space) were also discarded. The optimal box size of 486 pixels was selected based on the CTF aliasing frequency limit^22^. The optimal mask diameter of 136 Å was selected by executing multiple 3D refinement runs. To refine the per-particle defocus, beam tilt, and beam-induced motion corrections after the initial 3D refinement, the cycle of CTF refinement^44^ and Bayesian polishing^45^ in RELION3 was repeated three time. The number of repeats ensured that there were no further improvements. To measure the degree of improvement, 3D refinement (C3 symmetry, 136 Å mask diameter, no padding) with a soft-edged 3D mask (no pixel extension, 6-pixel soft cosine edge) and solvent-flattened FSCs options was used after each CTF refinement and Bayesian polishing step. After each 3D refinement, the gold-standard FSC resolution with a 0.143 criterion^24^ was used to estimate the global resolution using phase randomization to account for the possible artifactual resolution enhancement caused by the solvent mask^46^. To check the heterogeneity of the particle stack at this point, the 3D classification was conducted again by assuming asymmetry and 4 classes. Prior to this process, the 3D refinement after the second run of CTF refinement was re-executed assuming asymmetry instead of C3 symmetry to avoid the effect of an imposed symmetry bias in the classification. The second run was chosen because it had the highest resolution up to this point. Three 3D-classes showing clear high-resolution features were selected and yielded a stack of 121,353 particles. Subsequently, 3D refinement, Bayesian polishing, and then 3D refinement were conducted with the same settings as for the above refinement cycles imposing C3 symmetry again. The resolution of the final density map was 2.99 Å. The re-runs of the last Bayesian polishing excluding various numbers of the last movie frames to adjust the total electron exposure did not improve the reconstruction. The local resolution of the final reconstruction was estimated using the RELION3’s own implementation.

*Ac*NiR at pH 8.1 was processed similarly. For SPHIRE crYOLO particle picking, the generalized model with a box size of 128 pixels and selection threshold of 0.4 was used. The micrographs containing less than 30 picks were discarded. A stack of 176,256 particle images was extracted from 470 dose-weighted sum micrographs (3.3 Å/pixel, 64-pixel box size) and processed with RELION3 2D classification (200 expected classes, 120 Å mask diameter). The best 39 classes containing the 129,298 particles were used for RELION3 *ab initio* reconstruction (asymmetry, single expected class). The initial model was used for RELION3 3D classification (4 expected classes) after imposing C3 symmetry and low-pass filtering to 15 Å. The 3D class structures showed no significant differences, indicating high homogeneity of the dataset. The best 3D class was used as an initial model for the subsequent 3D refinement after rescaling to 0.88 Å/pixel with 486-pixel box size and low-pass filtering to 15 Å. Using the same rescale settings, 89,513 particles were also re-centered and re-extracted. The micrographs with defocus of >2.5 μm were removed prior to this extraction. Then, the particle images were 3D auto-refined without a 3D mask (C3 symmetry, 164 Å mask diameter, no padding) using RELION^3^. The 486-pixel box size was optimal according to the CTF aliasing frequency limit that ensures no CTF aliasing up to 2. 62 Å for 2.5 um defocus. Multiple runs of the 3D refinement indicated that the optimal mask diameter was 164 Å. After the initial 3D refinement, the three cycles of CTF refinement and Bayesian polishing in RELION3 were used. The third cycle did not yield any significant improvements. 3D refinement (C3 symmetry, 164 Å mask diameter, no padding) with a soft-edged 3D mask (no pixel extension, 6-pixel soft cosine edge) and solvent-flattened FSCs options was used after each step. The last Bayesian polishing was repeated by excluding the last 8 movie frames to adjust the total electron exposure from 50 to 42 e^−^/Å^2^. The final 3D refinement generated the 2.85 Å reconstruction. The local resolution of this density map was estimated with the RELION3’s own implementation.

For the visualization of the output 2D/3D images, UCFS Chimera^47^ and *e2display.py* of EMAN2^48^ were used.

### Equal-volume 3D difference

We devised the equal-volume 3D difference map method to examine the differences between the two cryo-EM density maps at the T2Cu site. The calculation of the difference between two independent cryo-EM density maps reconstructed from the independent datasets of different chemical environments is challenging because the exact scale factors of the density values are unknown, and these factors can be altered by various factors (e.g., different buffer conditions or average ice thicknesses). Therefore, to normalize the two maps, we used the equal-volume approach, since the compositions of the amino acid sequences are identical for *Ac*NiR at pH 6.2 and pH 8.1.

The equal-volume 3D difference map was calculated as follows. The procedure started from mutually aligning the local resolution filtered maps of the final reconstruction results of both datasets, using the “Fit in Map” function of the UCFS Chimera^47^. Using Moon Eliminator in SPHIRE^49^, the threshold of each density map was selected so that the two volumes occupied by the values above the thresholds were identical and represented the shapes of the side chains well. The selected thresholds were 0.0399 for pH 6.2 and 0.0405 for pH 8.1 (corresponding to the occupation of 80,614 voxels). Using the thresholds, both maps were binarized. At each voxel, the binary value of the pH 6.2 map was subtracted from the binary value of the pH 8.1 map, and finally, an equal-volume 3D difference map was generated. At each voxel in this map, the value −1 (blue blob in Fig. 5) indicates that the density exists only in the pH 6.2 structure at this 3D position, while the value 1 (red blob in Fig. 5) means the density exits only in the pH 8.1 structure. The value zero shows that both structures do or do not have the density.

The same binary normalization procedure was also used in part of the calculations of the 3D density existence probability map described below.

### 3D density existence probability map

The 3D density existence probability map was designed to show the quantitative significance of the difference between the cryo-EM and crystal structures of the *Ac*NiR, specifically in residues 185–215 (Fig. 2c, d), Ideally, the true variance of the density at each voxel should be calculated in the 3D space. Since each particle image reflects one single structure (or conformational state), it will be a simple task to calculate the 3D variance only if we could reconstruct the structure from the single image. However, this is impossible in reality. Therefore, we compromised by using the 3D variance of the averages of reconstructions from the mutually independent subsets. Here, each subset will consist of a certain number of particle images so that the 3D reconstruction is possible while maintaining a reasonable resolution for our purposes. Strictly speaking, the 3D variance of full-set reconstruction is not equivalent to the 3D variance of the averages of the subset reconstructions. However, the two variances will become closer as the number of subsets becomes larger, and they will eventually become equal when the subsets reach a single particle image (but this is not possible). Although the statistical bootstrap method could be used to select the subsets, we decided to use the 3D classification instead in order to emphasize the structural differences.

For the calculation of the 3D density existence probability map with each *Ac*NiR dataset, the following procedure was used. First, the final reconstruction result of each dataset was subjected to no-alignment 3D classification (asymmetry, no low-pass filtering) using RELION3, after expanding the particle stack by C3 symmetry using the relion_particle_symmetry_expand command^44^. The number of expected classes was set to 8 mainly by balancing among the validity of the statistic (larger is higher), the resolution of the 3D density existence probability map (larger is lower), and the processing time (larger is longer). The density maps of all 3D classes were binarized by the same procedure as used for the equal-volume 3D difference using the same density threshold value of each dataset. Then, the binarized maps of all 3D classes were added with the weighting based on the class distribution of the no-alignment 3D classification, and the 3D density existence probability map was obtained. In this map, a voxel value lower than 1/8 (i.e., the threshold value to render Fig. 2c, d) was considered to indicate no probability of any density existence at this 3D position at a 7/8 confidence level.

### Model building, refinement, and validation

The molecular structures of *Ac*NiR at pH 6.2 and 8.1 were refined against the maps obtained from the cryo-EM images. The PHENIX suite (version 1.14-3260)^50^ and Coot (version 0.8.9.2)^51^ were used during the refinements. The map quality analysis was carried out using *phenix.mtriage* before the refinement cycles. The symmetries of the maps were identified to be C3 by *phenix.map_symmetry*. A crystal structure of monomeric model *Ac*NiR (2bw4) was taken from PDB, and then all the alternative conformations and water/ligand molecules except copper ions were removed by *phenix.pdbtools*. This model was docked in the cryo-EM maps by *phenix.dock_in_map*, and the trimeric models were generated using the map symmetry information by *phenix.apply_ncs* and *phenix.auto_sharpen*. The refinement cycles were performed using *phenix.real_space_refinement* under the non-crystallographic symmetry (NCS) constraint, and visual model corrections were performed on Coot. The final models were validated by MolProbity^52^.

### Comparison of tertiary structures

The 3D structural comparison of *Ac*NiR molecules was performed using CA atoms from residues 10 to 330 with the program LSQKAB in the CCP4 program suite^53^. Coordinates of the crystal structures were obtained from PDB^54^. Comprehensive LSQ fittings were done by self-made PERL and shell scripts.

### EPR Spectroscopy

X-band EPR spectra were recorded by an RE-3X spectrometer (JEOL). The protein concentration of *Ac*NiR samples was 0.5 mM in 50 mM phosphate buffer (pH 6.0 and 8.0). The swept magnetic field was 300 ± 100 mT with a modulation amplitude of 1 mT. The sample temperature was maintained at 77 K. The g factors (g_‖_) and the hyperfine coupling constants (hfcc or A_‖_) of the type 1 and type 2 Cu sites of *Ac*NiR were obtained by the simulation.

## Data availability

The cryo-EM maps were deposited in the 3D-EM database (www.ebi.ac.uk/pdbe) with accession codes EMD-0730 and EMD-0731. The model coordinates were deposited in the PDB database with accession codes 6knf and 6kng.

## Acknowledgements

This work was supported by Grants-in-Aid from the Japanese Ministry of Education, Culture, Sports, Science, and Technology (nos. 17-214, 18-190 and 19-198 to T.K). This work was also supported by the Platform Project for Supporting Drug Discovery and Life Science Research (Basis for Supporting Innovative Drug Discovery and Life Science Research (BINDS)) from the Japan Agency for Medical Research and Development (AMED) under Grant Number JP20am0101071 (supporting nos. 1401 and 1630). The cryo-EM data were collected at the Cryo-EM facility in KEK (Ibaraki, Japan).

## Author contributions

N.A., T.Y., T.M., T.K., and T.S. conceived and designed the experiments. T.Y. purified the samples. N.A., T.Y., and M.K. prepared the cryo-EM samples and collected data. N.A., T.M., and M.K. processed the cryo-EM data. T.M., K.K., A.S., Y.Y., and F.Y. established the environment of data analysis. T.Y., T.K., and T.S. performed model building, refinement, and validation. N.A., T.Y., T.M., T.K., and T.S. prepared the manuscript.

## Additional information

Supplementary Information accompanies this paper at http://

Competing interests: The authors declare no competing financial interests.

Reprints and permission information is available online at http://

How to cite this article:

## Materials & Correspondence

All data needed to evaluate the conclusions in the paper are present in the paper and/or the Supplementary Materials. Additional data related to this paper may be requested from the authors. The cryo-EM maps are deposited in the Electron Microscopy Data Bank under accession codes EMD-0730 and EMD-0631. Structure coordinates are deposited at the PDB (PDB ID: 6knf and 6kng).

