## Supplementary Information for "Sub-3 Å resolution structure of 110 kDa nitrite reductase determined by 200 kV cryogenic electron microscopy"

**Supplementary Table 1. Statics of single-particle cryo-EM data and atomic model refinement**

| | NiR, pH6.2 | NiR, pH8.1 |
|---|---|---|
| ***Data collection*** | | |
| Microscope | FEI Talos Arctica | FEI Talos Arctica |
| Voltage [kV] | 200 | 200 |
| Detector | Falcon 3EC | Falcon 3EC |
| Magnification | 120 k | 120 k |
| Pixel size [Å/pixel] | 0.88 | 0.88 |
| Automation software | EPU | EPU |
| Total exposure [$e^-$/Å$^2$] | 50 | 50 |
| Exposure rate [$e^-$/Å$^2$ | 1.02 | 1.02 |
| Number of frames | 49 | 49 |
| Defocus range [µm] | 1.5 to 3 | 1 to 3 |
| Micrographs | 794 | 694 |
| Particles for Class2D | 285,529 | 176,256 |
| ***Reconstruction*** | | |
| Symmetry | C3 | C3 |
| Particles | 121,353 | 89,513 |
| Resolution [Å] | 2.99 | 2.85 |
| ***Model Refinement*** | | |
| Program | *phenix.real_space_refine* | *phenix.real_space_refine* |
| Resolution limit | 2.99 | 2.85 |
| Number of chains | 3 | 3 |
| No. of residues | 1002 | 1002 |
| No. of non-hydrogen | 7749 | 7749 |
| RMS bond length | 0.007(0) | 0.005(0) |
| RMS bond angle | 0.635(0) | 0.658(6) |
| Ramachandran plot | | |
| Preferred [%] | 96.39 | 99.10 |
| Allowed [%] | 3.61 | 0.90 |
| Outliers [%] | 0.00 | 0.00 |
| MolProbity score | | |
| Clash score | 7.65 | 5.30 |
| Rotamer outliers [%] | 8.82 | 3.68 |
| Overall score | 2.38 | 1.71 |
| EMDB/PDB codes | EMD-0730/6knf | EMD-0731/6kng |

**Supplementary Table 2. List of PDB IDs for *Ac*NiR stored in PDB**

| | |
|---|---|
| Group A | 1nia†, 1nib, 1nid, 1nif, 1nic, 1nie, 2nrd, 1kcb, 6gbb, 6qwg, 6gcg, 6gby, 6gtj, 6gb8, 5ogg, 5ofg, 1rzp, 5ofh, 5og3, 5off, 5og4, 5ogf, 5og2, 5og6, 5og5, 2bwd, 5i6l, 5i6n, 5i6k, 5i6m |
| Group B | 6gtn, 6gtl, 1rzq, 5n8f, 5n8g, 5n8i, 6gt2, 5n8h, 2bwi, 5i6o, 5akr, 5ofc, 5ofe, 5ofd, 2bw4, 5of8, 2bw5, 5i6p, 5of7, 6gti, 6gtk, 5of6, 5of5, 2y1a, 6gt0, 6gsq† |

† Representative crystal structures of each group used for further analysis.

**Supplementary Table 3. Statistics of pairwise comparisons of the *Ac*NiR's X-ray structures in PDB**

| | |
|---|---|
| **No. of PDBs** | 56 |
| **Highest resolution (Å)** | 0.8 |
| **Lowest resolution (Å)** | 2.7 |
| **Average of resolution** | 1.55 ± 0.39 |
| **No. of pairs** | 1,540 |
| **Minimum RMSD (Å)** | 0.0260 |
| **Maximum RMSD (Å)** | 0.7150 |
| **Average of RMSD (Å)** | 0.3563 |
| **SD of RMSD (Å)** | 0.1712 |

* LSQ fittings were performed using the trimer form of *Ac*NiR.

**Supplementary Table 4. Comparison of X-ray and cryo-EM structures of *Ac*NiR**

| | **Monomer** | | **Trimer** | |
|---|---|---|---|---|
| **pH** | **6.2** | **8.1** | **6.2** | **8.1** |
| **Minimum RMSD (Å)** | 0.434 | 0.436 | 0.584 | 0.588 |
| **Maximum RMSD (Å)** | 0.741 | 0.736 | 1.077 | 1.077 |
| **Average of RMSD (Å)** | 0.582 | 0.572 | 0.826 | 0.821 |
| **SD of RMSD (Å)** | 0.088 | 0.087 | 0.132 | 0.130 |

**Supplementary Table 5. EPR parameters of the type 1 and type 2 Cu sites**

| **pH** | **T1Cu** | | **T2Cu** | | | |
|---|---|---|---|---|---|---|
| | | | **Component 1** | | **Component 2** | |
| | $g_{\parallel}$ | $A_{\parallel}$ | $g_{\parallel}$ | $A_{\parallel}$ | $g_{\parallel}$ | $A_{\parallel}$ |
| **pH 6.0** | 2.18 | 7.2 | 2.34 | 13.1 | - | - |
| **pH 8.0** | 2.18 | 7.2 | 2.34 | 13.1 | 2.27 | 16.2 |

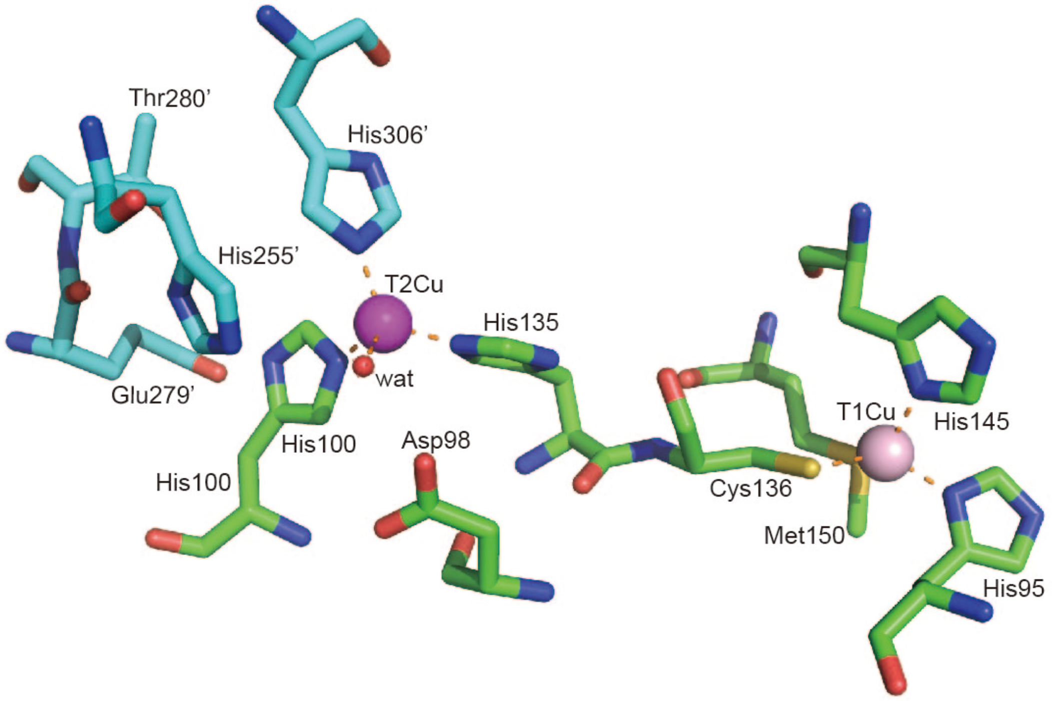


**Supplementary Fig. 1. Structure of AcNiR around the Cu sites**

Carbon atoms in subunits A and B are shown in green and cyan, respectively. T1Cu (pink), T2Cu (purple), and associated water molecule (red) are shown in sphere representations. The PDB ID of the coordinates is 1nia[9].

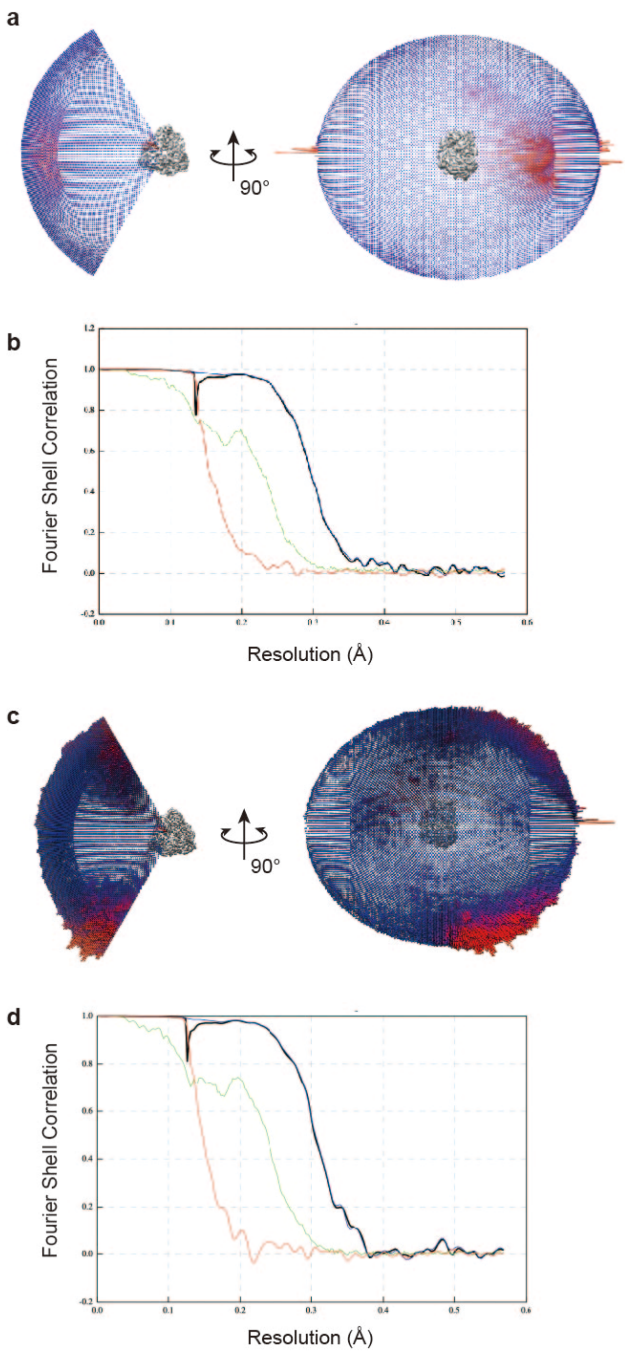


**Supplementary Fig. 2. Cryo-EM SPA validation of *Ac*NiR**

**a, b** Orientation distribution (**a**) and FSC curves (**b**) of the cryo-EM structure of *Ac*NiR at pH 6.2. **c, d** Orientation distribution (**c**) and FSC curves (**d**) of the cryo-EM structure of *Ac*NiR at pH 8.1.

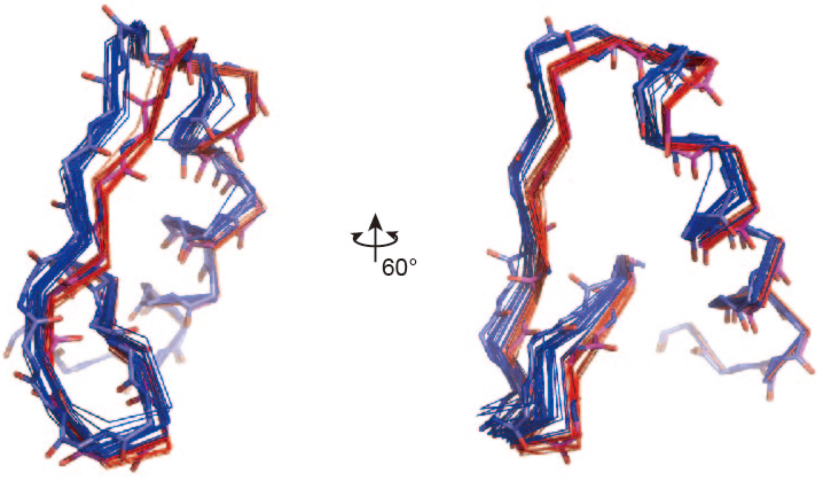


**Supplementary Fig. 3. Superposed structures of residues 185–215**

Residues 185–215 of the 56 *Ac*NiR crystal structures are superposed. Structures of groups A and B are shown in blue and red, respectively. Main chain structures of 1nia[9] and 6gsq[28] are shown in stick models.

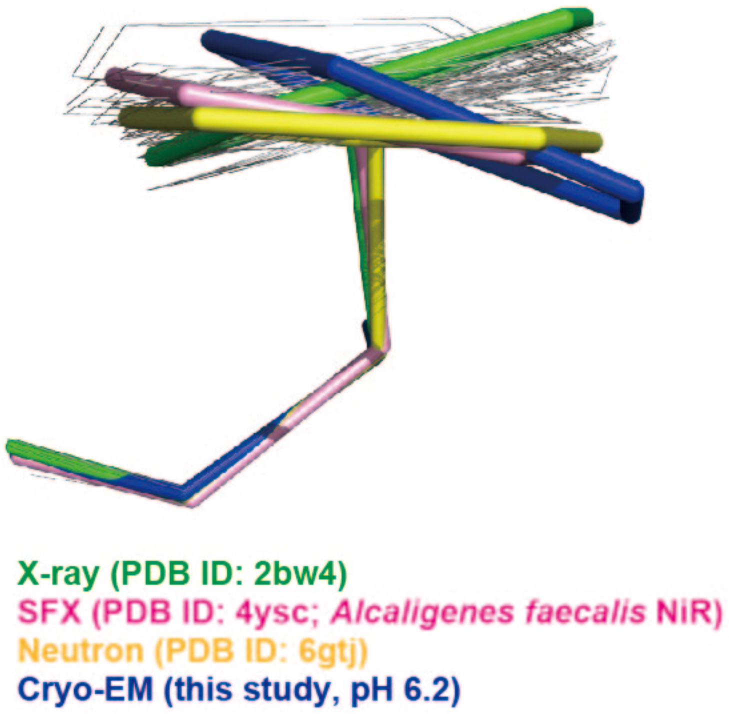


**Supplementary Fig. 4. Side-chain conformations of His255**

His255 regions of the 56 *Ac*NiR crystal structures (black lines) are superposed with neutron (PDB ID: 6gtj; yellow) and SFX (PDB ID: 4ysc; pink) structures. His255 of 2bw4 (green) adopts a typical structure of the reduced form. His255 of the cryo-EM structure (pH 6.2; blue) is in the oxidized form.

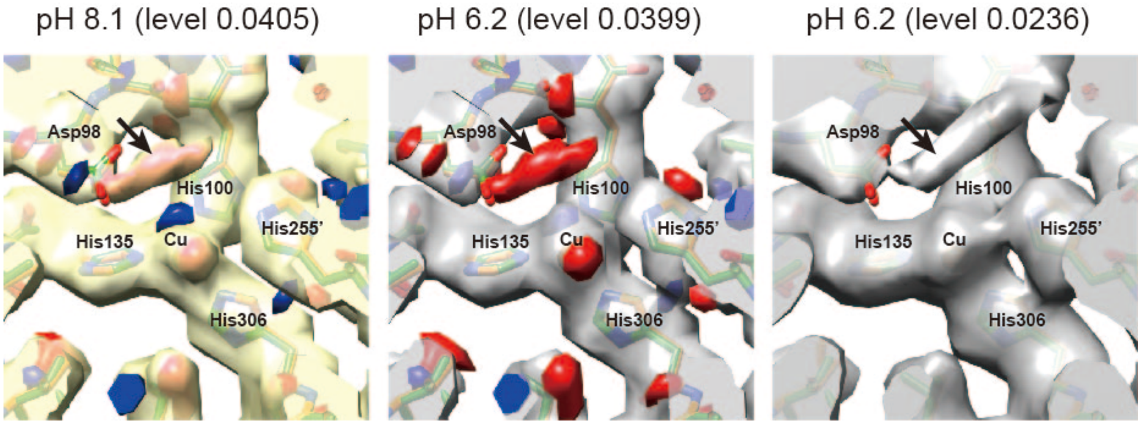


**Supplementary Fig. 5. Potential issues with the equal-volume 3D difference method**
A problematic large difference near Asp98 (the red blob indicated by an arrow in left and middle panels) of the difference map. The difference map is the same as in Fig. 5. The position of the large difference near Asp98 is overlapped with that of the bridging water molecule, which connects Asp98 and His255' (Adman et al., 1995). This indicates that the presumed bridging water disappears at pH 6.2, although the original pH 6.2 Cryo-EM map before the binarization shows the weak density of the presumed bridging water molecule with a threshold value lower than the one used for the binarization (arrow in the right panel). Since our interest was the structural change, the threshold value was selected to maintain clear shapes of the side chains. This criterion, along with the resolution difference between the pH 6.2 and pH 8.1 maps, likely caused the disappearance of the presumed bridging water from the binarized pH 6.2 map. Our approach in the equal-volume 3D difference map method seems to be valid, since it produced a result consistent with the previous studies and our EPR spectra (Fig. 6), indicating the directional change of the C-O bond of the main chain carbonyl oxygen of Glu279' (Fig. 5). However, the method seems to be sensitive to the threshold value used in the binarization step. Therefore, at the current level of resolution, this method should be considered more of a guide or support than a stand-alone proof. The interpretation of the difference blobs should be done carefully using other types of measurements. However, at higher resolution, the ambiguity of densities would be reduced, so the method would become less sensitive to the choice of binarization threshold. In the near future, the equal-volume 3D difference approach could become a reliable tool as the quality of Cryo-EM maps becomes sufficiently high.
